# Hopper: A Mathematically Optimal Algorithm for Sketching Biological Data

**DOI:** 10.1101/835033

**Authors:** Benjamin DeMeo, Bonnie Berger

## Abstract

Single-cell RNA-sequencing (scRNA-seq) has grown massively in scale since its inception, presenting substantial analytic and computational challenges. Even simple downstream analyses, such as dimensionality reduction and clustering, require days of runtime and hundreds of gigabytes of memory for today’s largest datasets. In addition, current methods often favor common cell types, and miss salient biological features captured by small cell populations. Here we present Hopper, a single-cell toolkit that both speeds up the analysis of single-cell datasets and highlights their transcriptional diversity by intelligent subsampling, or *sketching*. Hopper realizes the optimal polynomial-time approximation of the Hausdorff distance between the full and downsampled dataset, ensuring that each cell is well-represented by some cell in the sample. Unlike prior sketching methods, Hopper adds points iteratively and allows for additional sampling from regions of interest, enabling fast and targeted multi-resolution analyses. In a dataset of over 1.3 million mouse brain cells, we detect a cluster of just 64 macrophages expressing inflammatory tissues (0.004% of the full dataset) from a Hopper sketch containing just 5,000 cells, and several other small but biologically interesting immune cell populations invisible to analysis of the full data. On an even larger dataset consisting of ~2 million developing mouse organ cells, we show even representation of important cell types in small sketch sizes, in contrast with prior sketching methods. By condensing transcriptional information encoded in large datasets, Hopper grants the individual user with a laptop the same analytic capabilities as large consortium.

## 1 Introduction

Recent improvements in single cell technologies have enabled high-throughput profiling of individual cells, allowing fine-grained analyses of biological tissues. Droplet-based technologies have enabled profiling of millions of cells in a single experiment. Even larger datasets, containing tens or hundreds of millions or even billions of cells, are imminent [1]. For example, the Human Cell Atlas project aims to characterize and classify all cells in the human body [11].

While these large-scale assays have enormous scientific and therapeutic potential, they present significant computational and analytic challenges. Even the most basic exploratory analyses – visualization, clustering, and removal of batch effects – require time quadratic or worse in the size of the input data, which becomes intractable for more than tens of thousands of cells. Clinically or scientifically-relevant cells are often far outnumbered by common cell types [6]. Thus, there is a pressing need to produce *sketches* that reduce the size of single-cell datasets while preserving their transcriptional diversity.

There are several recent methods with this aim. Dropclust [13] performs Louvain clustering on an approximate nearest-neighbor network, and uses the resulting clusters as points of reference for downsampling. However, clustering itself is a very difficult and computationally expensive task, with the quality of the resulting sketches depending entirely on the clustering algorithm. The recently-introduced Geometric Sketching [6] samples evenly across transcriptional space by covering the PCA-reduced dataset with a gapped grid of disjoint axis-aligned hypercubes, and sampling a point at random from each. Geometric sketching is very fast, and outperforms DropClust in several key metrics [6]; yet, as we shall show, the fixed gridding axis can lead to artificial clusters near the grid intersections, potentially negatively effecting downstream analyses. Moreover, neither of these methods provides mathematical guarantees as to the mathematical approximation quality of the output sketches.

To address these challenges, we introduce Hopper, a novel toolkit that produces sketches with mathematical optimality guarantees on the distance from a point in the original data to the nearest point in the sketch. It achieves this result by implementing *farthest-first traversal*, a provably optimal polynomial-time approximation to the *k*-center problem. Intuitively, this means that every point in the full dataset *X* is very close to some point in the sketch *S*. Furthermore, unlike prior methods, Hopper allows fast insertion and removal of cells from the sketch, whilst preserving the strong mathematical guarantees. This enables fast multi-resolution analyses of large datasets.

While farthest-first traversal is mathematically powerful, its runtime is prohibitive for large sketches. We introduce two speedups using basic geometry which make the method feasible even on today’s largest datasets. To accommodate future datasets with tens or hundreds of millions of cells, Hopper implements a fully tunable pre-partitioning step, which reduces the runtime by orders of magnitude without significant loss in performance.

The code for Hopper is available at https://github.com/bendemeo/hopper. In addition, we have provided sketches of size 50,000 of the two largest single-cell datasets, available at http://cb.csail.mit.edu/cb/hopper/.

## 2 Results

### 2.1 Overview of Algorithm

At the core of Hopper is the *Farthest-first traversal*, an elegant greedy approximation to the *k*-center problem. Here to goal is to minimize, for some subset *S* of size *k* of a ground set *X*, the *Hausdorff distance*

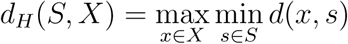

where *d* is a metric of choice (in our experiments, we use the Euclidean metric).

The algorithm works by sampling an initial point from *X* at random, and repeatedly adding the point *p* that is furthest from any of the previously-sampled points – that is,

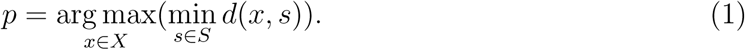

Intuitively, we repeatedly add to *S* the point of *X* that is least well-represented by *S*. We implement this in the `hop` function of the class `Hopper` in the Hopper module.

By design, this method is guaranteed to strictly decrease the Hausdorff distance *d*_*H*_ (*X, S*) after each step, assuming that the maximum is realized by only one point. In fact, one can show the following:

#### Theorem 1.

*Suppose that S is a k-step farthest traversal of X. Then*,

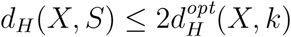

*where 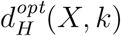 is the optimal Hausdorff distance realized by any subset of size k*.

Thus, farthest-first traversal realizes a 2-approximation to the optimal Hausdorff distance. The proof of this Theorem is found in [3]. The following theorem, due to [7], shows that we cannot reasonably hope to do any better:

#### Theorem 2.

*Let α <* 2. *Then, unless P* = *NP, there is no polynomial time algorithm for producing a set S satisfying*

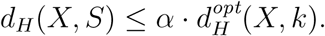

Thus, Hopper provides a gold-standard for sketching in the sense that no algorithm can reliably obtain a better Hausdorff distance, unless *P* = *NP*. The output of Hopper is an ordered collection of *x*_1_, *x*_2_, …, *x*_*k*_ of cells from *X*, such that for any ℓ ≤ *k*, the subset *x*_1_, …, *x*_ℓ_ reaches within a factor of two of the lowest possible Hausdorff distance for any sketch of size *ℓ*.

#### Geometric Speedups

The most computationally expensive aspect of farthest-first traversals is identifying the point *p* from equation 1. To do so, one must maintain for each *x ∈ X* the distance to the nearest point in *S*. Each time a point is added to *S*, these distances must be updated. A naïve approach computes the distance from every *x ∈ X* to the newly-added *p*, and updates the minimum distances accordingly. This requires *O*(*n*) time for each point addition, where *n* is the size of *X*. Producing a sketch of size *k* thus takes *O*(*nk*) time, which can be prohibitive for large sketches of large datasets.

Various speedups have been proposed in the theoretical computer science community (e.g. [5]), but all scale poorly with the dimensionality of the dataset. Instead, Hopper implements two simple geometric speedups using the triangle inequality. First, if the newly added point *p* has distance *r* to its nearest representative in *S*, then by the triangle inequality,

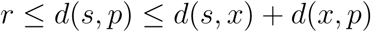

for any *s ∈ S* and *x ∈ X*. In particular, if *d*(*x, p*) ≤ *d*(*s, x*), then we must have 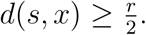. Thus, we need only examine those points in *X* with distance 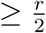 to their nearest point in *S*. To quickly find these points, the points *X* are sorted by their distance to the nearest point of *S*. Second, for *s ∈ S*, if *d*(*s, p*) ≥ 2*r*, then if *x ∈ X* is closest to *s*, the triangle inequality gives:

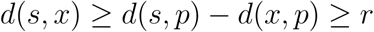

so there is no need to update any of the points associated to *s*. These two observations often allow significantly fewer than *n* points to be examined at each iteration. The exact runtime depends on the dimensionality and geometry of the dataset, but in practice the speedup is noticeable, especially for the first few thousand cells (Figure 2).

#### Pre-partitioning for even faster runtimes: the Treehopper class

As demonstrated by Geometric Sketching [6], spatial partitions can be used to very quickly generate sketches. The `Treehopper` class leverages this finding to further speed up sketch generation via *pre-partitioning*. The dataset *X* is first partitioned into subsets *X*_1_, …, *X*_*d*_, using a method of the user’s choice. A `Hopper` object *H*_*i*_ is instantiated in each partition *X*_*i*_, beginning a far traversal *S*_*i*_ of *X*_*i*_. These `Hoppers` are sorted according to their Hausdorff distance *d*_*H*_ (*S*_*i*_, *X*_*i*_). At each step, the `Hopper` with highest Hausdorff distance hops, adding a point to *S*_*i*_, and adjusting its position in the sorted list of `Hoppers`. The final sketch is the union of all the sub-traversals *S*_*i*_.

Using a fast heap implementation, this achieves an average hop time of 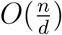 instead of *O*(*n*). Within each partition, the traversals *S*_*i*_ realize the optimality bound of Theorem 1, but this bound may not be achieved globally. This tradeoff between time and performance is fully tunable. If *d* = 1, we achieve optimal polynomial-time performance in *O*(*nk*) worst-case time. On the other extreme, if *d* = *n*, a random subsample is produced in *O*(*k*) time. For *d*-values in the tens to hundreds, these methods produce drastic speedups with little loss in accuracy (Figure 2)

In contrast with Geometric Sketching, where the partitions are all hypercubes of the same size and a point is drawn from each, `Treehopper` allows partitions to occupy variable-sized regions of transcriptional space, and draws variable numbers of points from the partitions according to their individual geometries. Thus, `Treehopper` bridges the gap between the very fast partition-and-sample approach and the slower, but mathematically optimal, farthest-first traversal approach. The choice of partition is entirely flexible; in our experiments, we use Principal Component Trees (PC-trees), which repeatedly split into equal halves along the leading principal component [14]. For the best possible performance, the partitions should be spatially separate and contain roughly equal numbers of points.

### 2.2 Experimental Results

#### Hopper better approximates biological datasets

We assessed our method’s performance on two of the largest published single-cell RNA-seq experiments: A set of 1.3 million mouse neurons from 10X genomics, and a set of *sim*2 million mammalian organogenesis cells [2]. Each dataset underwent standard normalization and feature selection protocols, and was projected to the its 100 Independent Components (ICs). Consistent with our mathematical guarantees, Hopper obtained Hausdorff distances significantly lower than any prior sketching technique, proving quantitatively that all cells in the dataset are better-represented (Figure 1). These improvements remained significant even when `Treehopper` was used with as many as 256 pre-partitions, suggesting that pre-partitioning does not substantially reduce performance. In contrast with Geometric Sketching, Hausdorff distance decreases smoothly as the number of points increases, likely because our methods are highly sensitive to individual outliers.

**Fig. 1:**
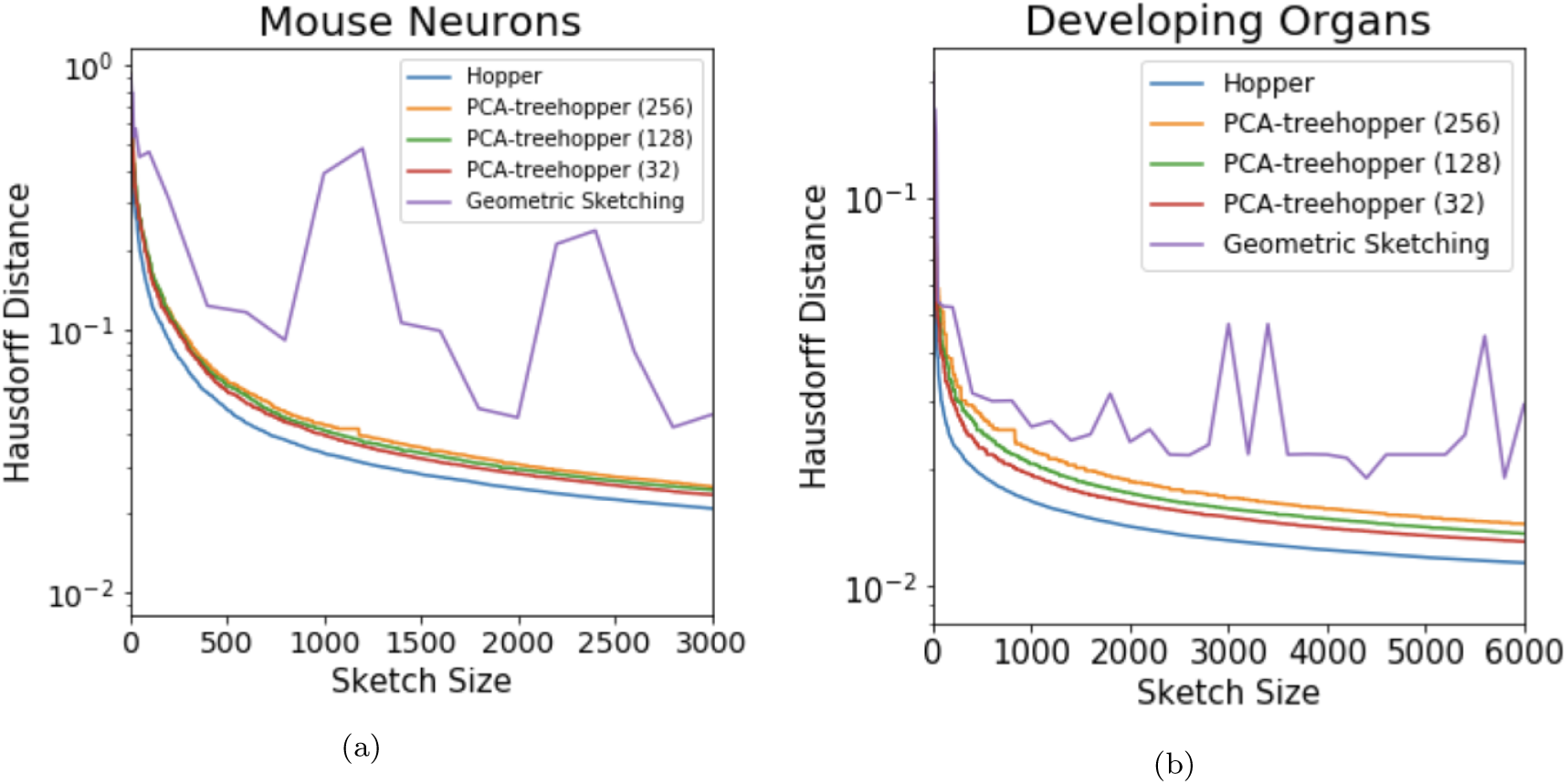
Hausdorff distances and runtimes for various Hopper routines, with Geometric Sketching for comparison, on (a) ~1.3 million mouse neurons and (b) ~2 million developing organ cells. The plain `Hopper` routine produces the lowest Hausdorff distance obtainable in polynomial time, with the faster `Treehopper` routines nearly realizing the optimum. All significantly outperform Geometric Sketching, and show more consistent Hausdorff performance.

As expected, `Hopper` and `Treehopper` run approximately linearly in the dataset size, with slopes depending on the number of pre-partitions (Figure 2). Geometric Sketching shows variable time performance between the two tested datasets. We suspect that because Geometric Sketching relies on a binary search to select the correct grid size, performance is heavily impacted by the number of search iterations needed, which is in turn influenced by the geometry of the data. Because even small sketch sizes may require several iterations, this leads to slower performance for small sketch sizes. On the other hand, the runtime is far less dependent on the sketch size, and may be faster for larger sketches (Figure 2b).

**Fig. 2:**
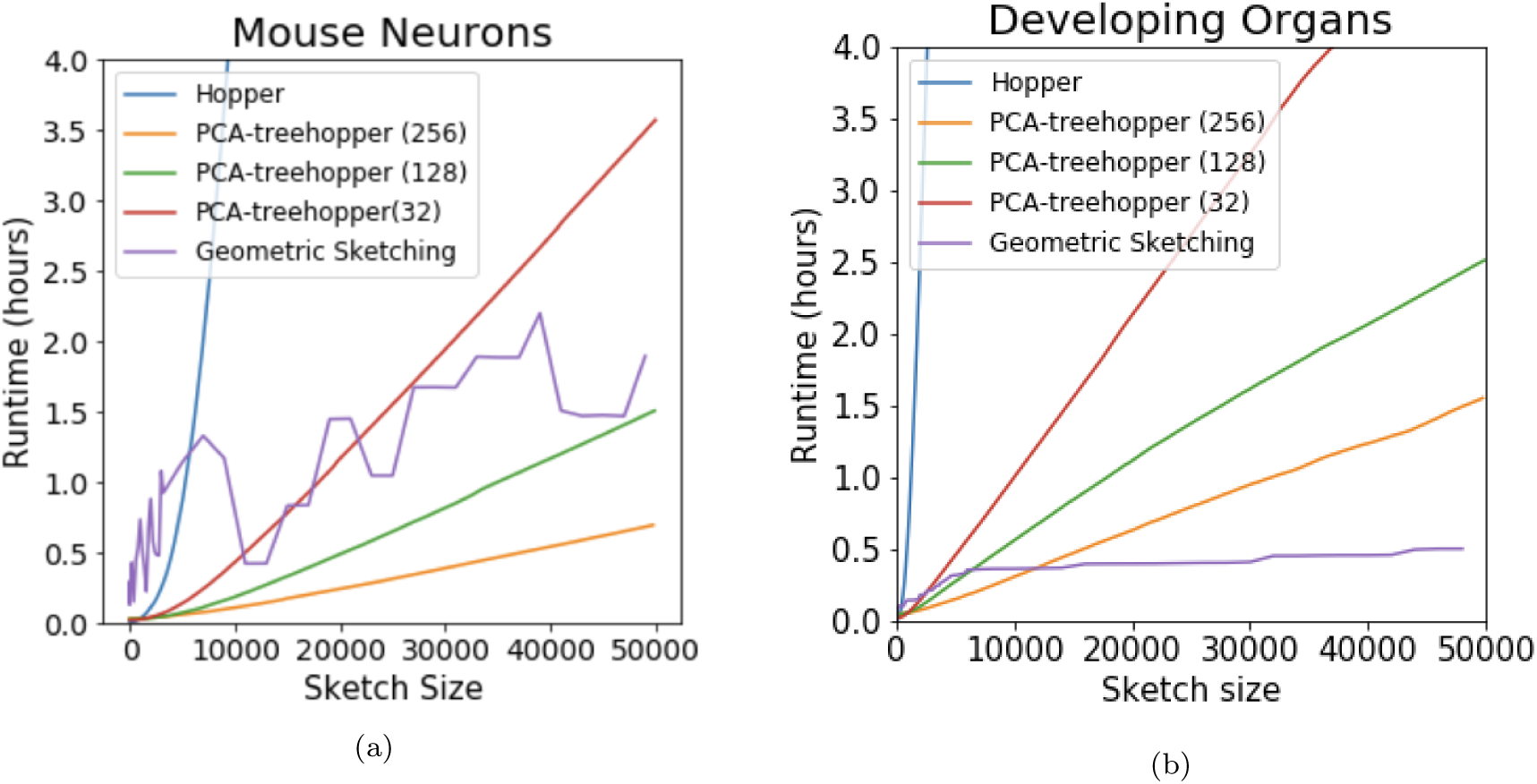
Runtimes for various sketching routines on the two tested datasets. All Hopper routines are linear in the sketch size, and add approximately one cell per 5 milliseconds, per thousand cells in each partition. Geometric sketching performs variably depending on the dataset’s geometry.

#### Hopper reveals novel clusters of immune cells in mouse brain data

Clustering is a key step in the analysis of single-cell data, allowing identification of known cell types, and discovery of new cell types, in a sample. Hopper facilitates this by representing rare clusters even with small sketches. To show this, we used Hopper to order the first 5,000 cells (about 0.4 %) of the 1.3 million neuron dataset, and clustered the resulting cells using Louvain community detection. These cluster labels were then propagated to the full dataset via nearest-neighbor classification. The detected clusters, plotted and annotated in Figure 3a, reveal several small but interesting cellular populations. For example, one of the clusters, consisting of a mere 64 cells, showed elevated expression of the *Cd5l* gene, which is expressed by macrophages in inflamed tissues [12] (Figure 3a). Another cluster consisted of just 114 cells with elevated expression of *Pf4* and *F13a1*, marker genes for activated platelets [10] (Figure 3a; Figure 3c). Another, consisting of just 221 cells, showed elevated expression the Interferon-*β* gene *Ifnb1*, expressed in fibroblasts and monocytes in response to viral infection [8]. Clusters 2-4 express canonical microglial markers, highlighting the transcriptional diversity of this group [4, 9]. Figure 3c shows expression heatmaps for each of these genes. Considering the role of the immune system in modulating disease states, these clusters are likely clinically important despite their small size.

**Fig. 3:**
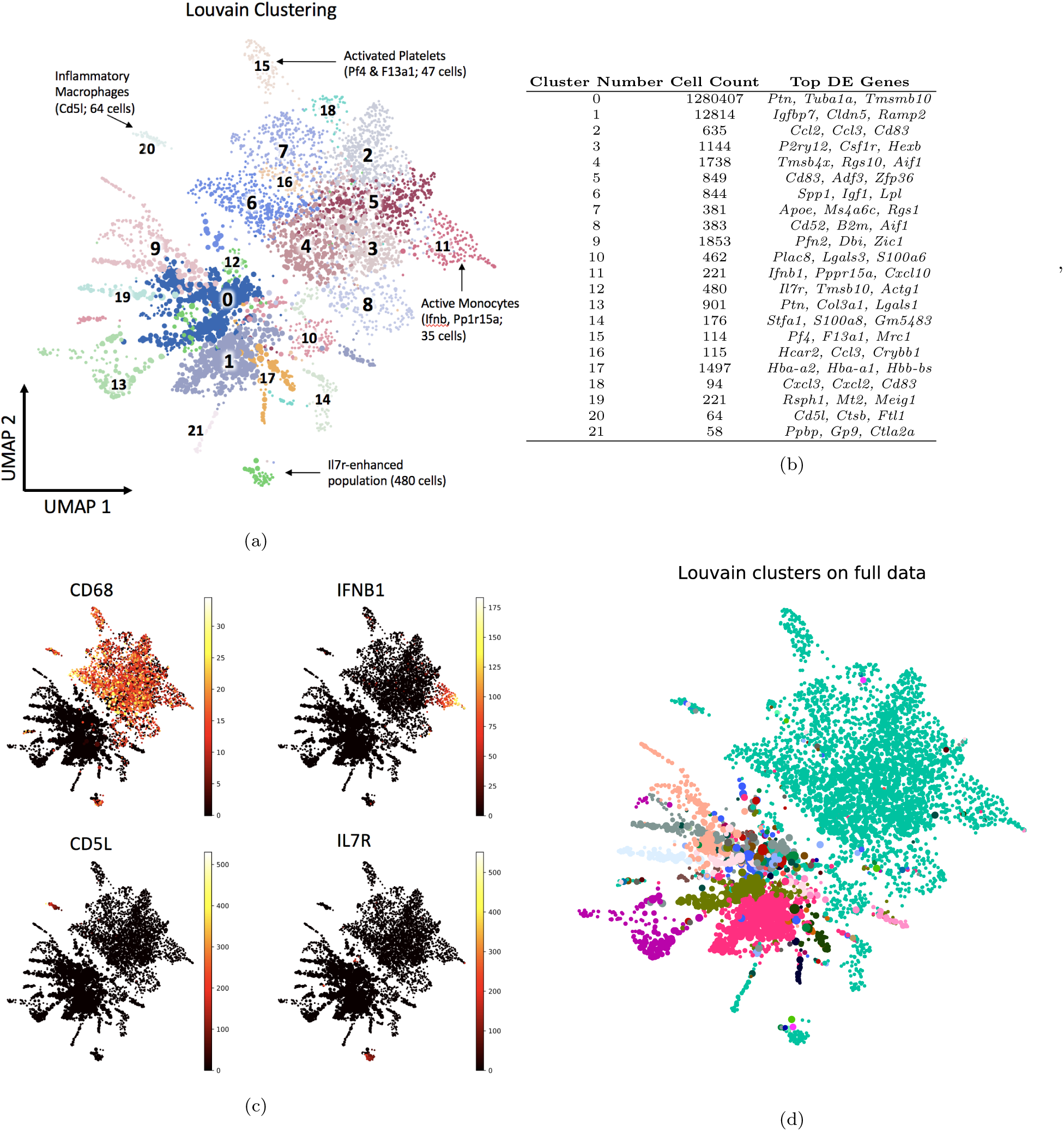
(a) Louvain clustering on the 5,000-point Hopper sketch of the 1.3 million-cell mouse brain dataset. Each cluster is numbered, and biologically interesting clusters are annotated with their inferred identity. (b) Table showing the cell counts per cluster after nearest-neighbor classification on the whole dataset, and the top differentially-expressed genes in each cluster. (c) Heat maps showing the expression of four different marker genes in a Hopper sketch of 5,000 mouse neurons out of 1.3 million. Elevated CD68 expression in the top half suggests a diverse population of immune cells. (d) Louvain clusters computed on the entire dataset fail to distinguish any of the cell subtypes identified by clustering on the sketch (see main text).

Table 3b lists all clusters, together with their sizes and differentially expressed genes relative to the total. Remarkably, almost all of the clusters computed from the Hopper sketch are extremely small relative to the full dataset size, indicating that miniscule populations can account for a large proportion of the dataset’s transcriptional diversity. These populations are completely invisible to any analysis of the full dataset. For example, the Louvain clustering produced by `scanpy` [16] on the full dataset lumps all of the immune cell clusters into a single relatively small cluster of 8,856 cells, obscuring their true diversity (Figure 3d).

#### Hopper samples smoothly across low-dimensional substructures

Geometric Sketching, the prior state of the art, covers the data with a gapped grid of axis-aligned boxes and samples a point from each box. This is a well-motivated approach that works very well on many datasets. However, we have observed that axis-aligned grid hypercubes do not always represent the data evenly, especially where the local low-dimensional structure of the data aligns poorly with the gridding axis. As demonstrated schematically in Figure 4a, this results in more points near the grid square intersections, an effect which is compounded with an increase in the ambient dimension *D*, since as many as 2^*D*^ hypercubes may meet. As a result, we observe clumping even when the underlying data is Gaussian (Figure 4b). On the mouse organogenesis dataset, this manifests as additional clusters not present in the Hopper sketches (Figure 4d).

**Fig. 4:**
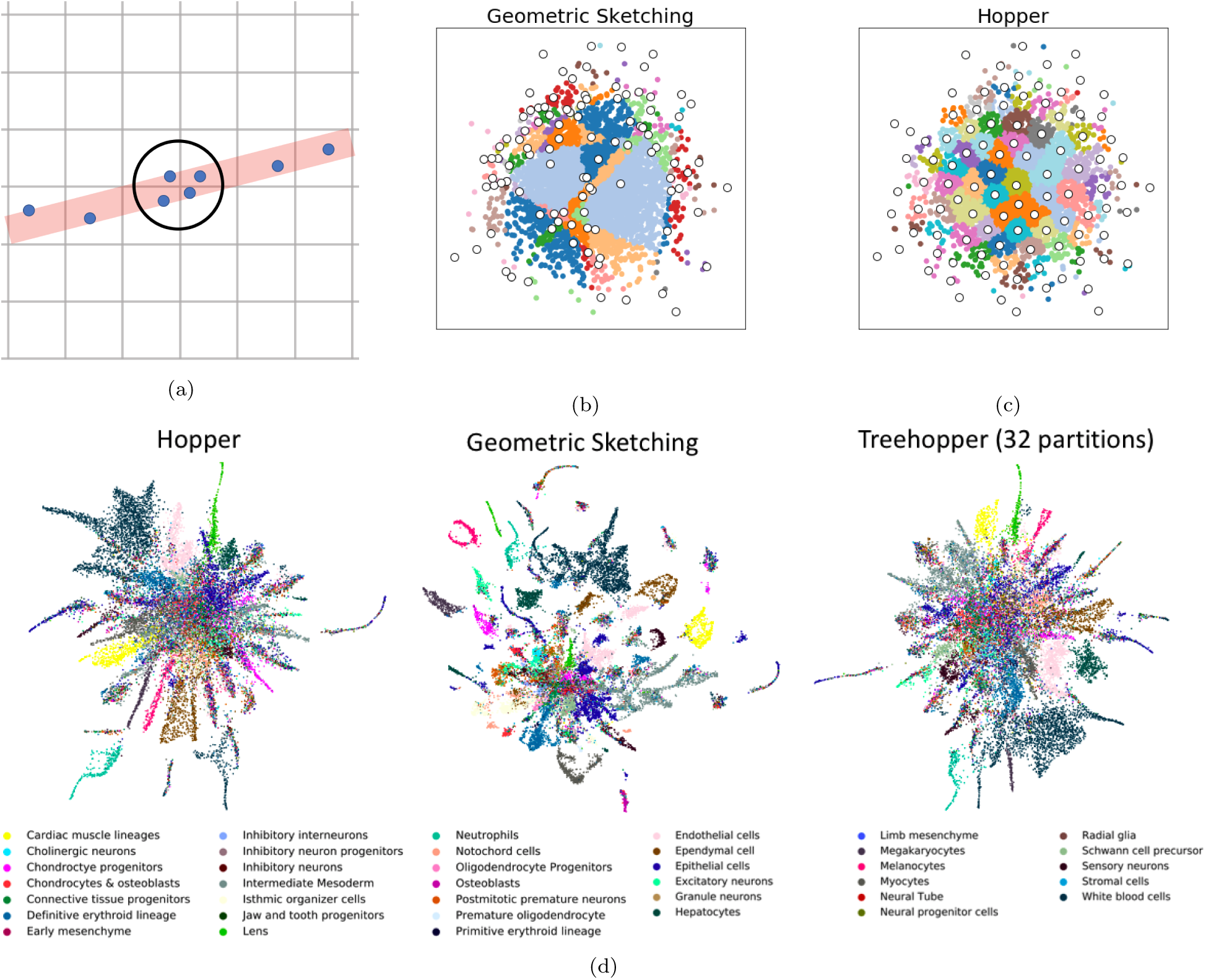
Grid-based sketches clump at grid intersections. (a) Schematic diagram, assuming the data lies near a one-dimensional line (red) in two-dimensional space. Where the line meets the grid intersection, four points are sampled, causing an artificial clump (circled). This effect is compounded in higher dimensions. (b) A sample geometric sketch on 2-D Gaussian data randomly embedded into 100-dimensional space. The 100 sampled points are shown in white, with the remaining points colored by grid cell. The grids partition the data erratically, and regions near grid intersections are preferentially sampled. (c) Hopper sketch of the same data, with 100 points colored according to their closest sampled point. The data is smoothly represented. (d) UMAP visualizations of sketches produced by `Hopper`, Geometric Sketching, and by `Treehopper` with 32 partitions, colored by cell type. Geometric Sketching generates additional clusters at grid intersections. Hopper and Treehopper avoid this issue.

Hopper avoids this issue entirely by not relying on any axis, ensuring that all low-dimensional substructures are smoothly represented regardless of spatial orientation (Figure 4c and Figure 4d). Sketches produced with Treehopper using PCA-trees to pre-partition closely resemble those of Hopper, even with partition sizes less than 5% of the total sample size (Figure 4d).

We note that the pure-partitioning approach taken by Geometric Sketching does allow remarkably fast runtimes, and the artificial clumping effect does not occur on all datasets; indeed, geometric sketches of the 1.3 million mouse neuron dataset closely resemble Hopper sketches (data not shown). We suspect that the observed defect emerges only when the data has high intrinsic dimensionality, i.e. lies on a high-dimensional manifold, because this allows for more high-dimensional intersections between occupied grid hypercubes.

## 3 Discussion

Hopper leverages the mathematical power of farthest-first traversal to produce sketches that preserve a sample’s transcriptional diversity and biological meaning. These sketches are mathematically guaranteed to represent the original data as well as any polynomial-time algorithm, thus providing a much-needed gold standard. By incorporating the powerful partition-and-sample approach, it allows tunable scaling to massive-scale single-cell datasets without excessive computational burden.

We have provided the first 50,000 cells in the far traversal of two super-massive single-cell RNA-seq datasets. This data requires only a few megabytes of storage, but allows immediate production of mathematically optimal sketches of any size smaller than 50,000. This allows the researcher immediate access both to small sketches, which may distill out the rare cell types, and larger sketches, which may be more comprehensive at the expense of obscuring rare cell types. Indeed, the position of a cell in the far traversal produced by Hopper may prove a valuable input to other downstream analyses. For example, one could modify the Louvain community detection algorithm by weighting vertices according to their traversal positions, and modifying the modularity-detection step to ensure that both rare and common clusters are represented.

The experiments in this paper exclusively use Euclidean distance as a measure of dissimilarity, but the Hopper framework generalizes to any dissimilarity measure. Unlike other methods, an explicit embedding of the cells is not required - only a method of determining distance. As demonstrated by kernel SVM, this is a highly desirable property. There are several existing algorithms for learning discriminative metrics from single cell datasets, which can be directly fed into the Hopper framework. For example, SIMLR [15] uses machine learning to jointly predict the clustering and the distance measure. Other possibilities abound, from established kernels (e.g. polynomial kernels or radial basis functions) to custom-designed kernels which may incorporate prior knowledge about the relevant factors shaping a dataset’s diversity. Because the distance function can be user-specified, inputting such custom kernels into the Hopper framework is very straightforward. While we expect most useful kernels to obey the triangle inequality, and while the fastest version of Hopper assumes that it holds, Hopper also accommodates arbitrary kernels via a small change in input parameters.

Hopper offers a flexible, scalable, and mathematically principled workflow for distilling the essence of a single-cell dataset. As these datasets grow larger, such methods will become increasingly vital for enabling the advanced and computationally-expensive downstream workflows that the future of single-cell data undoubtedly holds.

## Acknowledgments

The authors are grateful to Brian Hie, Hoon Cho, Ashwin Narayan, Rohit Singh, and other members of the Berger lab for valuable feedback and input. We also thank Bryan Bryson for his expert help in biologically interpreting the Louvain clusters of the mouse brain data, outlined in Figure 3.

